# Physiological and biochemical changes associated with acute experimental dehydration in the desert adapted mouse, *Peromyscus eremicus*

**DOI:** 10.1101/047704

**Authors:** Lauren Kordonowy, Kaelina D. Lombardo, Hannah L. Green, Molly Dawson, Evice A. Bolton, Sarah LaCourse, Matthew D. MacManes

**Affiliations:** University of New Hampshire, Department of Molecular Cellular and Biomedical Sciences; University of New Hampshire, Department of Biological Sciences; University of New Hampshire, Department of Psychology

**Keywords:** Dehydration, desert, electrolyte, rodent

## Abstract

Characterizing traits critical for adaptation to a given environment is an important first step in understanding how phenotypes evolve. How animals adapt to the extreme heat and aridity commonplace to deserts represents is an exceptionally interesting example of these processes, and has been the focus of study for decades. In contrast to those studies, where experiments are conducted on either wild animals or captive animals held in non-desert conditions, the study described here leverages a unique environmental chamber that replicates desert conditions for captive *Peromyscus eremicus* (cactus mouse). Here we establish baseline values for daily water intake and for serum electrolytes, as well as the response of these variables to experimental dehydration. In brief, *P. eremicus*’ daily water intake is very low. It’s serum electrolytes are distinct from many previously studied animals, and its response to acute dehydration is profound, though not suggestive of renal impairment, which is atypical of mammals.

**Summary statement:** The establishment of baseline values for serum electrolytes and water intake, as well as their response to acute dehydration is critical for characterizing the physiology necessary for desert survival.

**Conflict of Interest Statement:** The authors declare no conflict of interest.

## Introduction

Understanding the evolution of adaptive traits has long been one of the primary goals in evolutionary biology. The study of the relationships between fitness and phenotype, often powered by modern genomic techniques (59), has provided researchers with insight into the mechanistic processes that underlie adaptive phenotypes (15, 28). Systems in which the power of genomics can be combined with an understanding of natural history and physiology are well suited for the study of adaptation (9, 44) especially when researchers have the ability to assay the link between genotype and phenotype in wild animals and then conduct complementary experiments using representative animals in carefully controlled laboratory environments. The study described here, characterizing the physiology and serum biochemistry of *Peromyscus eremicus* is the first step in a larger study aimed at understanding the genomics architecture of adaptation to desert environments.

Desert adaptation has significant ecological, evolutionary, and biomedical significance. In contrast to humans and other mammals, desert rodents can survive in extreme environmental conditions and are resistant to the effects of dehydration. Physiological adaptions to deserts have been characterized in several rodents. Specifically, renal histology has been studied in multiple Heteromyid rodents (3), and the general conclusion is that these desert adapted animals have evolved elongate Loops of Henle (7, 10, 38) that are hypothesized to optimize water conservation. In addition to studies of renal histology, several studies have characterized pulmonary water loss (23, 51), water metabolism (26), and water consumption (12, 34, 35, 41, 46) in desert rodents. While desert animals possess specialized physiology that is efficient with regards to water metabolism and loss, whether or not specialized genomic adaptation exists is an active area of research (32, 36, 37).

Although the cactus mouse (*Peromyscus eremicus*) has not been a particular focus for the study of desert adaptation (but see (2, 32), this Cricetid rodent native to the arid regions of the Southwestern United States and Northern Mexico (57) offers a unique opportunity to understand physiological adaptations to deserts. *P. eremicus* is a member of a larger genus of animals known colloquially as the “*Drosophila* of mammals” (9), and *Peromyscus* species have been the focus of extensive study (25, 33, 52, 54). *P. eremicus* is a sister species to the non-desert adapted *P. californicus* (13), and it is closely related to *P. crinitus*, the canyon mouse, which is another desert adapted rodent native to Southwestern deserts.

Critical to desert survival is the ability to maintain water balance even when the acute loss of water exceeds dietary water intake (24). Indeed, the mammalian corpus consists of 60% water (30). Far from a static reservoir, proper physiologic function requires water for numerous processes, including nutrient transport (22), signal transduction, pH balance, thermal regulation (42) and the removal of metabolic waste. To accomplish these functions, a nearly constant supply of water is required to replace water loss (30), which occurs mainly via the gastrointestinal and genitourinary systems, and evaporative loss, which is greatly accelerated in extreme heat and aridity (16). Because the body possesses limited reserves, when loss exceeds intake during even a short period of time, dehydration and death can occur. Mammals are exquisitely sensitive to dehydration and possess limited compensatory mechanisms.

Characterizing desert adaptation requires careful and integrative physiological studies, which should include a detailed characterization of water intake, responses to dehydration, and the measurement of blood electrolytes. Indeed, quantifying these metrics is one of the first steps in understanding how animals survive in the extreme heat and aridity of deserts. In particular, the electrolytes chloride and sodium are important markers of dehydration (18). These molecules play essential roles in metabolic and physiological processes, and they are integral to the functionally of a variety of transmembrane transport pumps (11, 29), neurotransmission (62), and maintenance of tonicity (19). Furthermore, hypernatremia causes restlessness, lethargy, muscle weakness, or coma (1). Bicarbonate ion, in contrast, is primarily responsible for aiding in the maintenance of the acid-base balance and is resorbed in the renal tubules (39). Blood urea nitrogen (BUN) is a test that assays the abundance of urea – the end-product for metabolism of nitrogen containing compounds. Urea is resorbed in the glomerulus, and renal impairment is often inferred when BUN becomes elevated (8). Importantly, the canonical model of urea resorption is dependent on urine volume, which is markedly diminished in desert rodents, thus limiting the utility of using BUN as an indicator of renal function. Lastly, creatinine, a product of muscle breakdown, whose measured level does not depend on urine volume is used as a measure of renal function (8).

Genes most frequently implicated in desert-adaptation include members of the aquaporin family (27). However, previous work suggests that an alternative gene family, the solute carriers, are more relevant for desert-adaptation in the cactus mouse (32). As a first step towards fully elucidating the patterns of adaptive evolution to deserts in *P. eremicus*, we characterized the normal patterns of water intake and electrolyte levels as well as the physiologic response to experimental dehydration. As such, this study provides critical physiological and biochemical information about *P. eremicus* and its response to dehydration and is generally useful as researchers begin to leverage large-scale genome data against classic questions regarding the evolution of adaptive phenotypes.

## Materials and Methods

We used captive *P. eremicus* (n=44, 24 male, 20 female) that were descendant from mice purchased from the University of South Carolina Peromyscus Genetic Stock Center. The USC colony was founded using wild caught animals from a dry-desert population in Arizona. For ongoing experimental purposes, animals are housed in a large walk-in environmental chamber built to replicate the environmental conditions in which this population has evolved. Specifically, the animals experience a normal diurnal pattern of temperature fluctuation, ranging from 90F during the daytime to 75F during the night. Relative humidity (RH) ranges from 10% during the day to 25% during the night. Animals are housed in standard lab mouse cages with bedding that has been dehydrated to match desert conditions. They are fed a standard rodent chow, which has also been dehydrated. Water is provided *ad lib* during certain phases of experimentation and withheld completely during others. All animal care procedures follow the guidelines established by the American Society of Mammalogy (53) and have been approved by the University of New Hampshire Animal Care and Use Committee under protocol number 103092.

All animals included in this study were sexually mature adults, as defined for males as having scrotal testes and for females as having a perforate vaginal meatus. A slight bias for the inclusion of males exists, as a concurrent study of male reproductive genomics was occurring. Preliminary analyses conducted suggest that no significant differences in any of the physiological measures, and thus, males and females were analyzed as one group. For a subset of animals, water intake was measured, which was accomplished via the use of customized 15ml conical tubes, wherein water intake was measured every 24 hours for a minimum of 3 consecutive days (range 3-10 days). Animals selected for the dehydration trial were weighed on a digital scale, housed without water for three days, then re-weighed to determine the change in body mass due to dehydration. At the conclusion of water measurement or after a three-day dehydration animals were sacrificed via isoflurane overdose and decapitation. Immediately after death, a 120uL sample of trunk blood was obtained for serum electrolyte measurement. This was accomplished using an Abaxis Vetscan VS2 machine with a critical care cartridge, which measures the concentration of several electrolytes (Sodium, Chloride, Bicarbonate ion, Creatinine, and Blood Urea Nitrogen (BUN)) relevant to hydration status and renal function. Lastly, the kidney, spleen, liver, lung, hypothalamus, testes, vas deferens and epididymis were dissected out and stored in RNAlater (Ambion Inc.) for future study. All statistical analyses were carried out in the statistical package, R (50).

## Results

We measured the daily water intake for 44 adult cactus mice (24 male, 20 female) for between three and ten consecutive days. Mean water intake was 0.11 mL per day per gram body weight (median=0.11, SD=0.05, min=0.033, max=0.23). We measured levels of serum Sodium, Chloride, Bicarbonate ion, Creatinine, and Blood Urea Nitrogen (BUN) for the same 44 adult mice, thereby establishing normal (baseline) values for *P. eremicus* (Figure 1 and Table 1).

**Table 1.**
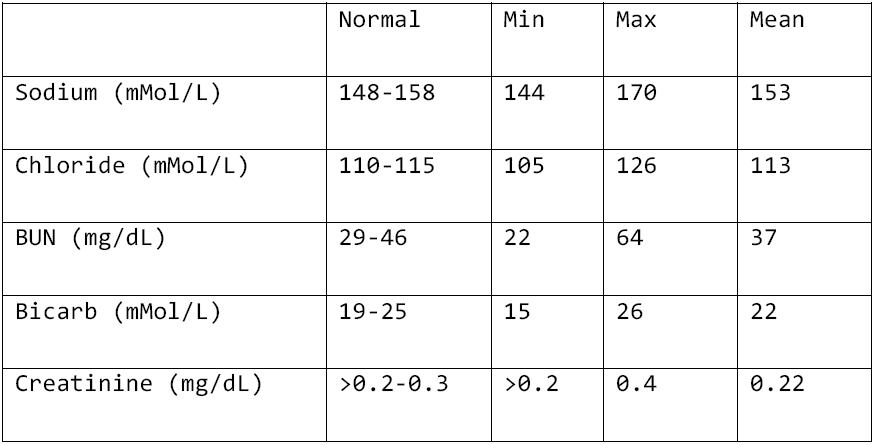
Normal values for serum electrolytes. Normal values (n=44, 24 male, 20 female) are defined as those values falling between the 1^st^ and 3^rd^ quartile. Of note, the Abaxis VS2 electrolyte analyzer does not measure Creatinine below 0.2 mg/dL; therefore, the range for normal Creatinine is truncated at this value.

**Figure 1.**
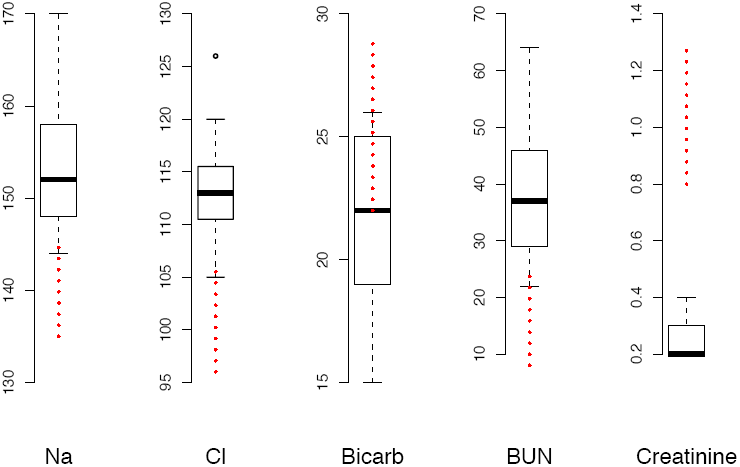
Normal values (n=44, 24 male, 20 female) for serum electrolytes. Human normal values (from Medline) are plotted for comparison in dotted red lines. Of note, the Abaxis VS2 electrolyte analyzer does not measure Creatinine below 0.2 mg/dL, and therefore the range for normal Creatinine is truncated at this value.

A comparison of mice provided with water *ad libitum* to mice that exposed to experimental water deprivation for three days revealed that the dehydrated mice lost an average of 23.2% of their body weight (median=23.9%, SD=5.3%, min=12.3%, max=32.3%, n=13 dehydration treatment, 7 males, 6 females). Despite this substantial weight loss, anecdotally, mice appeared healthy. They were active, eating, and interacting with handlers and other mice, normally. The amount of weight loss did not depend on daily water intake (p=0.63, R^2^= 0.03), though the trend suggests that animals that drink more water lost more weight). Furthermore, body weight did not strongly influence the percent loss of body weight (Figure 2; p=0.68, R^2^= 0.02).

**Figure 2.**
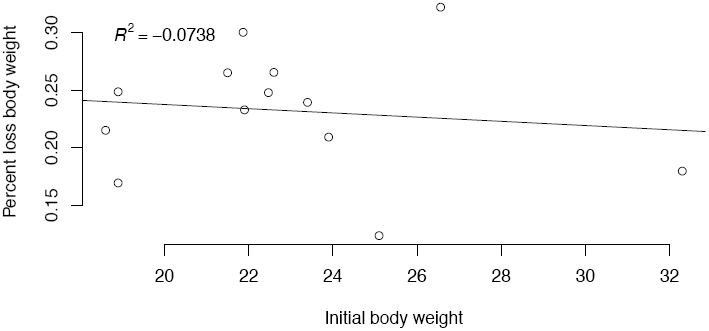
Percent body weight loss as a function of initial body weight due to experimental dehydration. No significant trend exists. (n=13, 7 males, 6 females)

In addition to a substantial loss in body weight, dehydration was associated with differences in serum electrolytes (Figure 3; n=13 dehydrated, n=31 hydrated). These changes were subtle, but significant using a two-sample t-test (p < 0.008 in all cases).

**Figure 3.**
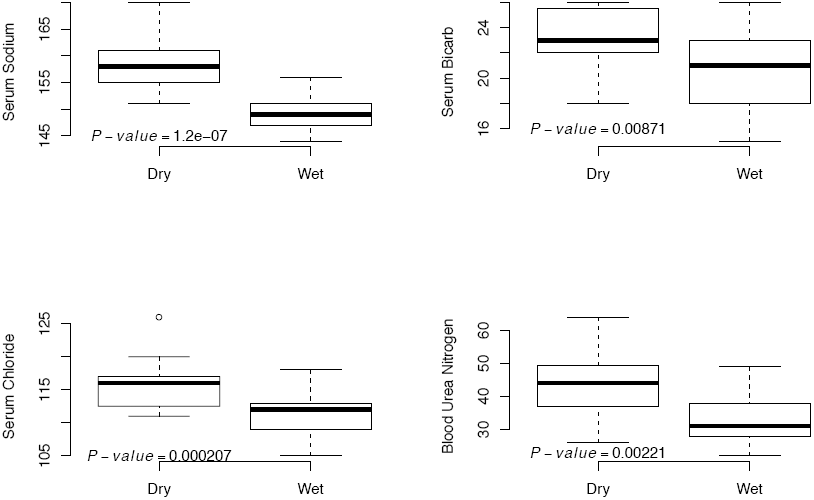
Experimental dehydration resulted in increases in serum sodium, chloride, BUN and bicarbonate ion. Reported p-values are from a two-tailed t-test (n=13 dehydrated: DRY, n=31 hydrated: Wet)

Lastly, the levels of serum electrolytes were tightly correlated with percent body weight loss (Figure 4). Indeed, the relationship between the level of serum sodium and weight loss was positive and significant, (ANOVA, F-statistic: 12.85, 11 DF, p= 0.004), as was the relationship between BUN and weight loss (ANOVA, F-statistic: 9.089, 11 DF, p= 0.012). The relationships between weight loss and chloride and bicarbonate levels respectively, were positive but not significant. Of note, all data will be deposited in Dryad upon acceptance for publication.

**Figure 4.**
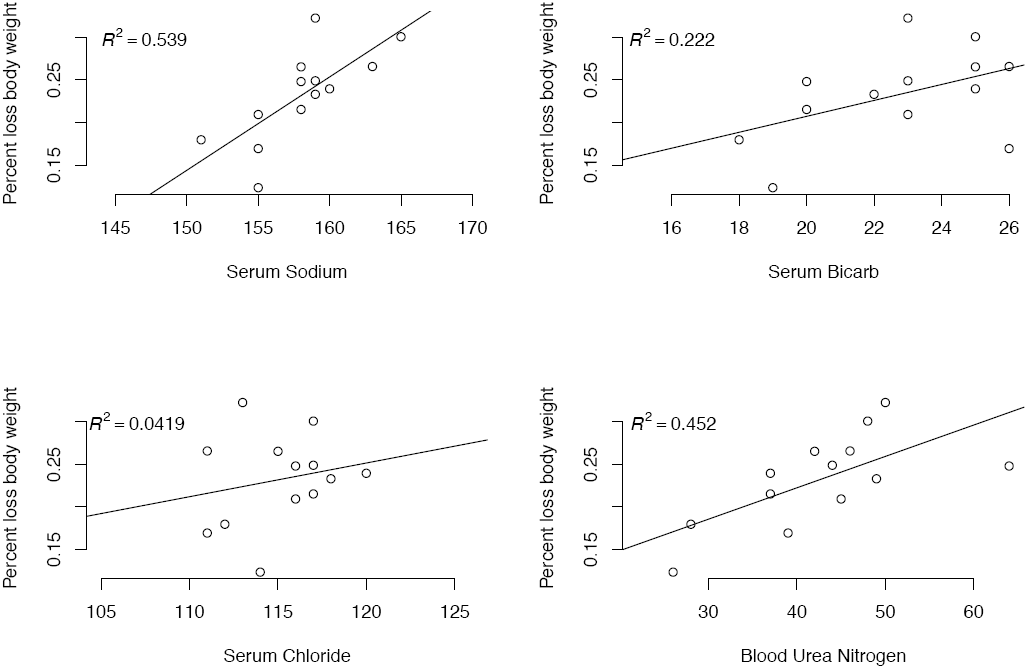
The relationship between serum electrolytes is positive in all cases and significant for Sodium (F-statistic: 12.85, 11 DF, p-value: 0.004283) and BUN (F-statistic: 9.089, 11 DF, p-value: 0.01177). n=13, 7 males, 6 females

## Discussion

Deserts are amongst the harshest environments on the planet. Indeed, animals living in these areas must be highly adapted to the unique combination of extreme heat and aridity. Given that our understanding of the physiology of desert adapted animals is limited largely to studies in renal histology (38) and on water intake and output (34, 56), an enhanced understanding of serum electrolyte changes due to dehydration is informative. Because many of the harmful effects of dehydration result from electrolyte abnormalities, characterizing normal values and the electrolyte response to dehydration represents a critical first step in garnering a deeper understanding of how desert animals survive despite severe and prolonged dehydration.

In this study, normal (baseline) values for serum Sodium, Chloride, Bicarbonate Ion, Creatinine, and Blood Urea Nitrogen were established in a captive colony of lab animals housed in desert conditions. Although these measures may differ in wild animals (see (14) for a brief review of such differences), establishing normal values in captive animals is crucial, though future studies aim to understand the patterns of electrolyte variation in wild animals. In *P. eremicus*, we define the normal ranges for each electrolyte as those values falling between the 1^st^ and 3^rd^ quartile. Serum Chloride and Sodium were significantly higher than in published ranges for other mammals, including humans, a marsupial (58), *Cricetomys* (48), and the porcupine (43). However, serum chloride and sodium levels in our study were quite comparable to another wild rodent, *Neotoma fuscipes* (61), a Mustelid (55), and the Hyrax (5). Values for BUN are generally higher in this study; unfortunately, a direct comparison is not possible, as measured values are dependent on the volume of urine produced. Serum Creatinine is low, largely resulting from the general lack of muscle mass in *P. eremicus* relative to other mammals. However, because the equipment used to analyze this electrolyte does not effectively capture the lower end of the biological range, direct comparisons are not made for this metric. Future measurements, using a more sensitive HPLC method for quantitating serum Creatinine will improve our ability to detect more subtle changed of change.

In addition to characterizing baseline electrolytes and their response to experimental dehydration, the normative value for daily water intake was estimated to be 0.11 mL per day per gram body weight. Though comparable measures of water consumption are scarce, one study in two arid adapted *Limoys* (*L. pictus* and *L. irroratus*) housed in non-desert captive settings were estimated to be 0.18 and 0.17 mL per day per gram body weight respectively (17) – a value much greater than in *P. eremicus*.

Animals that were exposed to experimental dehydration lost a substantial amount of body weight. Dehydration in humans, resulting in loss of even a fraction of this amount results in cardiovascular collapse and death (40). Indeed, even a dehydration-related loss of a few percent of body weight may cause serious renal impairment or renal failure. That the cactus mouse may lose so much weight as a result of dehydration and remain active, and apparently healthy, without renal impairment is a testament to their desert adaptation. The magnitude of weight loss and the negative (though non-significant) relationship between baseline weight and weight loss, coupled with the lack of behavioral impairments suggests that metabolic water production via the oxidation of fat may be an important and potentially adaptive mechanism preventing more serious complications from acute dehydration. Indeed, water metabolism may produce a substantial amount of water (reviewed in (31)), and had been demonstrated in a diverse group of animals including marine mammals (49) and desert rodents (20). Future studies of fat metabolism in *P. eremicus*, using computed tomography, are planned. Because animals are essentially anuric, particularly when dehydrated, means direct measurement of fax oxidation (*e.g.*, urine beta hydroxybutryate) is not possible.

As described above, mice appear grossly behaviorally intact. Despite this, they may be experiencing a degree of cognitive impairment, as is the case with human dehydration, where even mild-dehydration is associated with cognitive impairment (4). Future studies, using classical Y-maze and novel object recognition tests aim to understand more fully the cognitive effects of dehydration in cactus mouse.

In addition to weight loss, dehydrated animals demonstrated biochemical evidence of physiological stress, in the form of increased Sodium, Chloride, BUN, and Bicarb. There were no significant relationships between any physiological value and Creatinine, suggesting that dehydration related stress does not result in renal impairment or damage. Indeed, this is in contrast to humans and other mammals where acute dehydration of the nature imposed on these animals is universally related to renal failure and subsequent death. That *P. eremicus* can withstand this level of dehydration is a testament to the processes involved in adaptation.

In summary, we present here a set of physiological measurements that represent the endpoints in the physiological management of acute dehydration in the desert adapted cactus mouse. How these endpoint are achieved is an outstanding question deserving future study, particularly in light of global climate change (31). For instance Vasopressin, along with the Renin-Angiotensin-Aldosterone system are thought to be a critically important to the regulation of water and solute balance (6, 45, 47, 60, 63). Comparative genomic analysis, studies of gene expression, and the measurement of protein levels will provide important insights into the actual mechanisms underlying these phenotypes.

In addition to understanding the mechanisms of salt and water balance, characterizing the ways in which desert animals prevent dehydration-linked renal failure is exceptionally important. Unlike humans, where repeated dehydration events leads to a progressive decline in renal function (21), it is hypothesized that repeated acute-dehydration is unlikely to be linked to renal failure in animals that have evolved in desert environments. Testing this hypothesis, along with understanding the mechanisms which limit renal damage (*e.g.,* modulating renal microcirculation, maintaining cell volume via organic osmolytes) could provide previously uncharacterized clinically-relevant insights into renal (dis)function.

## References

1. Adrogué HJ, Madias NE. Hypernatremia. New Engl J Med 342: 1493–1499, 2000.

2. al-Kahtani MA, Zuleta C, Caviedes-Vidal E, Garland T. Kidney mass and relative medullary thickness of rodents in relation to habitat, body size, and phylogeny. Physiol Biochem Zool 77: 346–365, 2004.

3. Altschuler EM, Nagle RB, Braun EJ, Lindstedt SL, Krutzsch PH. Morphological study of the desert heteromyid kidney with emphasis on the genus *Perognathus*. Anat Rec 194: 461–468, 1979.

4. Armstrong LE, Ganio MS, Casa DJ, Lee EC, McDermott BP, Klau JF, Jimenez L, Le Bellego L, Chevillotte E, Lieberman HR. Mild dehydration affects mood in healthy young women. Journal of Nutrition 142: 382–388, 2012.

5. Aroch I, King R, Baneth G. Hematology and serum biochemistry values of trapped, healthy, free-ranging rock hyraxes (*Procavia capensis*) and their association with age, sex, and gestational status. Vet Clin Pathol 36: 40–48, 2007.

6. Bankir L, Fernandes S, Bardoux P, Bouby N, Bichet DG. Vasopressin-V2 receptor stimulation reduces sodium excretion in healthy humans. J Am Soc Nephrol 16: 1920–1928, 2005.

7. Barrett JM, Kriz W, Kaissling B, De Rouffignac C. The ultrastructure of the nephrons of the desert rodent (*Psammomys obesus*) kidney. II. Thin limbs of Henle of long-looped nephrons. Am J Anat 151: 499–514, 1978.

8. Baum N, Dichoso CC, Carlton CE. Blood urea nitrogen and serum creatinine. Physiology and interpretations. Urology 5: 583–588, 1975.

9. Bedford NL, Hoekstra HE. Peromyscus mice as a model for studying natural variation. eLife 4:e06813, 2015.

10. Beuchat CA. Structure and concentrating ability of the mammalian kidney: correlations with habitat. Am J Physiol 271: R157–79, 1996.

11. Blaustein MP, Lederer WJ. Sodium/calcium exchange: its physiological implications. Physiol Rev 79: 763–854, 1999.

12. Bradford DF. Water stress of free-living *Peromyscus truei*. Ecology 55: 1407–1414, 1974.

13. Bradley RD, Durish ND, Rogers DS, Miller JR, Engstrom MD, Kilpatrick CW. Toward a molecular phylogeny for *Peromyscus*: Evidence from mitochondrial cytochrome-b sequences. J Mammal 88: 1146–1159, 2007.

14. Calisi RM, Bentley GE. Lab and field experiments: Are they the same animal? Horm Behav 56: 1–10, 2009.

15. Castoe TA, de Koning APJ, Hall KT, Card DC, Schield DR, Fujita MK, Ruggiero RP, Degner JF, Daza JM, Gu W, Reyes-Velasco J, Shaney KJ, Castoe JM, Fox SE, Poole AW, Polanco D, Dobry J, Vandewege MW, Li Q, Schott RK, Kapusta A, Minx P, Feschotte C, Uetz P, Ray DA, Hoffmann FG, Bogden R, Smith EN, Chang BSW, Vonk FJ, Casewell NR, Henkel CV, Richardson MK, Mackessy SP, Bronikowsi AM, Yandell M, Warren WC, Secor SM, Pollock DD. The Burmese python genome reveals the molecular basis for extreme adaptation in snakes. PNAS 110: 20645–20650, 2013.

16. Cheuvront SN, Kenefick RW, Montain SJ, Sawka MN. Mechanisms of aerobic performance impairment with heat stress and dehydration. Journal of Applied Physiology 109: 1989–1995, 2010.

17. Christian DP, Matson JO, Rosenberg SG. Comparative water balance in two species of *Liomys*. Comp Biochem Physiol, Part A Mol Integr Physiol 61A: 589–559, 1978.

18. Costill DL, Coté R, Fink W. Muscle water and electrolytes following varied levels of dehydration in man. Journal of Applied Physiology 40: 6–11, 1976.

19. Feig PU, McCurdy DK. The hypertonic state. N Engl J Med 297: 1444–1454, 1977.

20. Frank CL. Diet Selection by a Heteromyid Rodent: Role of Net Metabolic Water Production. Ecology 69: 1943–1951, 1988.

21. Glaser J, Lemery J, Rajagopalan B, Diaz HF, García-Trabanino R, Taduri G, Madero M, Amarasinghe M, Abraham G, Anutrakulchai S, Jha V, Stenvinkel P, Roncal-Jimenez C, Lanaspa MA, Correa-Rotter R, Sheikh-Hamad D, Burdmann EA, Andres-Hernando A, Milagres T, Weiss I, Kanbay M, Wesseling C, Sánchez-Lozada LG, Johnson RJ. Climate Change and the Emergent Epidemic of CKD from Heat Stress in Rural Communities: The Case for Heat Stress Nephropathy. Clinical Journal of the American Society of Nephrology 11: 1472–1483, 2016.

22. Haussinger D. The role of cellular hydration in the regulation of cell function. Biochem J 313 ( Pt 3): 697–710, 1996.

23. Hayes J, Bible C, Boone J. Repeatability of mammalian physiology: Evaporative water loss and oxygen consumption of *Dipodomys merriami*. J Mammal 79: 475–485, 1998.

24. Heimeier RA, Davis BJ, Donald JA. The effect of water deprivation on the expression of atrial natriuretic peptide and its receptors in the spinifex hopping mouse, *Notomys alexis*. Comparative Biochemistry and Physiology Part A: Physiology 132: 893–903, 2002.

25. Hoekstra H, Moghadam HK, Hoekstra J, Harrison PW, Berrigan D, Zachar G, Székely T, Vignieri S, Mank JE, Hoang A, Hill C, Beerli P, Kingsolver J. Strength and tempo of directional selection in the wild. P Natl Acad Sci Usa 98: 9157–9160, 2001.

26. Howell AB, Gersh I. Conservation of water by the rodent *Dipodomys*. J Mammal 16: 1, 1935.

27. Huang Y, Tracy R, Walsberg GE, Makkinje A, Fang P, Brown D, Van Hoek AN. Absence of aquaporin-4 water channels from kidneys of the desert rodent *Dipodomys merriami merriami*. Am J Physiol-Renal 280: F794–F802, 2001.

28. Huerta-Sánchez E, Jin X, Asan, Bianba Z, Peter BM, Vinckenbosch N, Liang Y, Yi X, He M, Somel M, Ni P, Wang B, Ou X, Huasang, Luosang J, Cuo ZXP, Li K, Gao G, Yin Y, Wang W, Zhang X, Xu X, Yang H, Li Y, Wang J, Wang J, Nielsen R. Altitude adaptation in Tibetans caused by introgression of Denisovan-like DNA. Nature 512: 194–197, 2014.

29. Jentsch TJ, Stein V, Weinreich F, Zdebik AA. Molecular Structure and Physiological Function of Chloride Channels. Physiol Rev 82: 503–568, 2002.

30. Jéquier E, Constant F. Water as an essential nutrient: the physiological basis of hydration. Eur J Clin Nutr 64: 115–123, 2009.

31. Johnson RJ, Johnson RJ, Stenvinkel P, Stenvinkel P, Jensen T, Jensen T, Lanaspa MA, Lanaspa MA, Roncal C, Roncal C, Song Z, Song Z, Bankir L, Bankir L, Sánchez-Lozada LG, Sanchez-Lozada LG. Metabolic and Kidney Diseases in the Setting of Climate Change, Water Shortage, and Survival Factors. Journal of the American Society of Nephrology 27: 2247–2256, 2016.

32. MacManes MD, Eisen MB. Characterization of the transcriptome, nucleotide sequence polymorphism, and natural selection in the desert adapted mouse *Peromyscus eremicus*. PeerJ 2: e642, 2014.

33. MacManes MD, Lacey EA. Is promiscuity associated with enhanced selection on MHC-DQα in mice (genus *Peromyscus*)? PLOS ONE 7: e37562, 2012.

34. MacMillen RE, Lee AK. Australian desert mice: independence of exogenous water. Science 158: 383–385, 1967.

35. Mares M. Water Economy and Salt Balance in a South-American Desert Rodent, *Eligmodontia typus*. Comp Biochem Phys A 56: 325–332, 1977.

36. Marra NJ, Eo SH, Hale MC, Waser PM, DeWoody A. A priori and a posteriori approaches for finding genes of evolutionary interest in non-model species: Osmoregulatory genes in the kidney transcriptome of the desert rodent *Dipodomys spectabilis* (Banner-Tailed Kangaroo Rat). Comparative Biochemistry and Physiology - Part D: Genomics and Proteomics (July 26, 2012). doi: 10.1016/j.cbd.2012.07.001.

37. Marra NJ, Romero A, DeWoody JA. Natural selection and the genetic basis of osmoregulation in heteromyid rodents as revealed by RNA-seq. Mol Ecol 23: 2699–2711, 2014.

38. Mbassa GK. Mammalian renal modifications in dry environments. Vet Res Commun 12: 1–18, 1988.

39. McKinney TD, Burg MB. Bicarbonate and fluid absorption by renal proximal straight tubules. Kidney Int 12: 1–8, 1977.

40. Mehta RL, Cerdá J, Burdmann EA, Tonelli M, García-García G, Jha V, Susantitaphong P, Rocco M, Vanholder R, Sever MS, Cruz D, Jaber B, Lameire NH, Lombardi R, Lewington A, Feehally J, Finkelstein F, Levin N, Pannu N, Thomas B, Aronoff-Spencer E, Remuzzi G. International Society of Nephrology's 0by25 initiative for acute kidney injury (zero preventable deaths by 2025): a human rights case for nephrology. Lancet 385: 2616–2643, 2015.

41. Merkt JR, Taylor CR. “Metabolic switch” for desert survival. P Natl Acad Sci Usa 91: 12313–12316, 1994.

42. Montain S, Latzka W, Sawka N. Fluid replacement recommendations for training in hot weather. Mil Med 164: 502–508, 1999.

43. Moreau B, Vié JC, Cotellon P, Thoisy ID, Motard A, Raccurt CP. HEMATOLOGIC AND SERUM BIOCHEMISTRY VALUES IN TWO SPECIES OF FREE-RANGING PORCUPINES (*COENDOU PREHENSILIS, COENDOU MELANURUS*) IN FRENCH GUIANA. J Zoo Wildl Med 34: 159–162, 2003.

44. Mullen LM, Vignieri SN, Gore JA, Hoekstra HE. Adaptive basis of geographic variation: genetic, phenotypic and environmental differences among beach mouse populations. Proc Biol Sci 276: 3809–3818, 2009.

45. Muñoz A, Riber C, Trigo P. Dehydration, electrolyte imbalances and renin-angiotensin-aldosterone-vasopressin axis in successful and unsuccessful endurance horses Equine Veterinary Journal - Wiley Online Library. Equine Veterinary … (2010). doi: 10.1111/j.2042-3306.2010.00211.x/pdf.

46. Nagy KA. Seasonal patterns of water and energy balance in desert vertebrates. J Arid Environ 14: 201–210, 1988.

47. Nielsen S, Chou C, Marples D, Christensen E, Kishore B, Knepper M. Vasopressin increases water permeability of kidney collecting duct by inducing translocation of aquaporin-CD water channels to plasma-membrane. P Natl Acad Sci Usa 92: 1013–1017, 1995.

48. Nssien M, Olayemi FO, Onwuka SK, Olusola A. Comparison of some plasma biochemcial parameters in two generations of african giant rat (*Cricetomys gambianus*, waterhouse). African Journal of Biomedical Research 5, 2002.

49. Ortiz RM. Osmoregulation in marine mammals. Journal of Experimental Biology 204: 1831–1844, 2001.

50. R Core Development Team F. R: A Language and Environment for Statistical Computing [Online]. R Development Core Team: 2011. http://www.R-project.org.

51. Schmidt-Nielsen B, Schmidt-Nielsen K. Pulmonary Water Loss in Desert Rodents. Am J Physiol 162: 31–36, 1950.

52. Shorter KR, Crossland JP, Webb D, Szalai G, Felder MR, Vrana PB. *Peromyscus* as a Mammalian Epigenetic Model. Genetics Research International 2012: 1–11, 2012.

53. Sikes RS, Gannon WL, Animal Care and Use Committee of the American Society of Mammalogists. Guidelines of the American Society of Mammalogists for the use of wild mammals in research. J Mammal 92: 235–253, 2011.

54. Steiner CC, Weber JN, Hoekstra HE. Adaptive variation in beach mice produced by two interacting pigmentation genes. PLoS Biol 5: e219, 2007.

55. Thornton PC, Wright PA, Sacra PJ, Goodier TE. The ferret, *Mustela putorius furo*, as a new species in toxicology. Lab Anim 13: 119–124, 1979.

56. Tracy R, Walsberg G. Intraspecific variation in water loss in a desert rodent, *Dipodomys merriami*. Ecology 82: 1130–1137, 2001.

57. Veal R, Caire W. Peromyscus eremicus. Mammalian Species 118: 1–6, 2001.

58. Viggers KL, Lindenmayer DB. Variation in Hematological and Serum Biochemical Values of the Mountain Brushtail Possum, *Trichosurus caninus* Ogilby (Marsupialia: Phalangeridae). J Wildlife Dis 32: 142–146, 1996.

59. Vignieri SN, Larson JG, Hoekstra HE. The selective advantage of crypsis in mice. Evolution 64: 2153–2158, 2010.

60. Weaver D, Walker L, Alcorn D, Skinner S. The contributions of renin and vasopressin to the adaptation of the Australian spinifex hopping mouse (*Notomys alexis*) to free water deprivation. Comp Biochem Physio 108: 107–116, 1994.

61. Weber DK, Danielson K, Wright S, Foley JE. Hematology and serum biochemistry values of dusky-footed wood rat (/). J Wildlife Dis 38: 576–582, 2002.

62. Yu FH, Catterall WA. Overview of the voltage-gated sodium channel family. Genome Biology 4: 207, 2003.

63. Zuber AM, Singer D, Penninger JM, Rossier BC, Firsov D. Increased renal responsiveness to vasopressin and enhanced V2 receptor signaling in RGS2-/- mice. J Am Soc Nephrol 18: 1672–1678, 2007.

